## Supplementary Figures 1-11 for "Chondrogenic Enhancer Landscape of Limb and Axial Skeleton Development"

### Supplementary Info

### Supplementary Figure 1

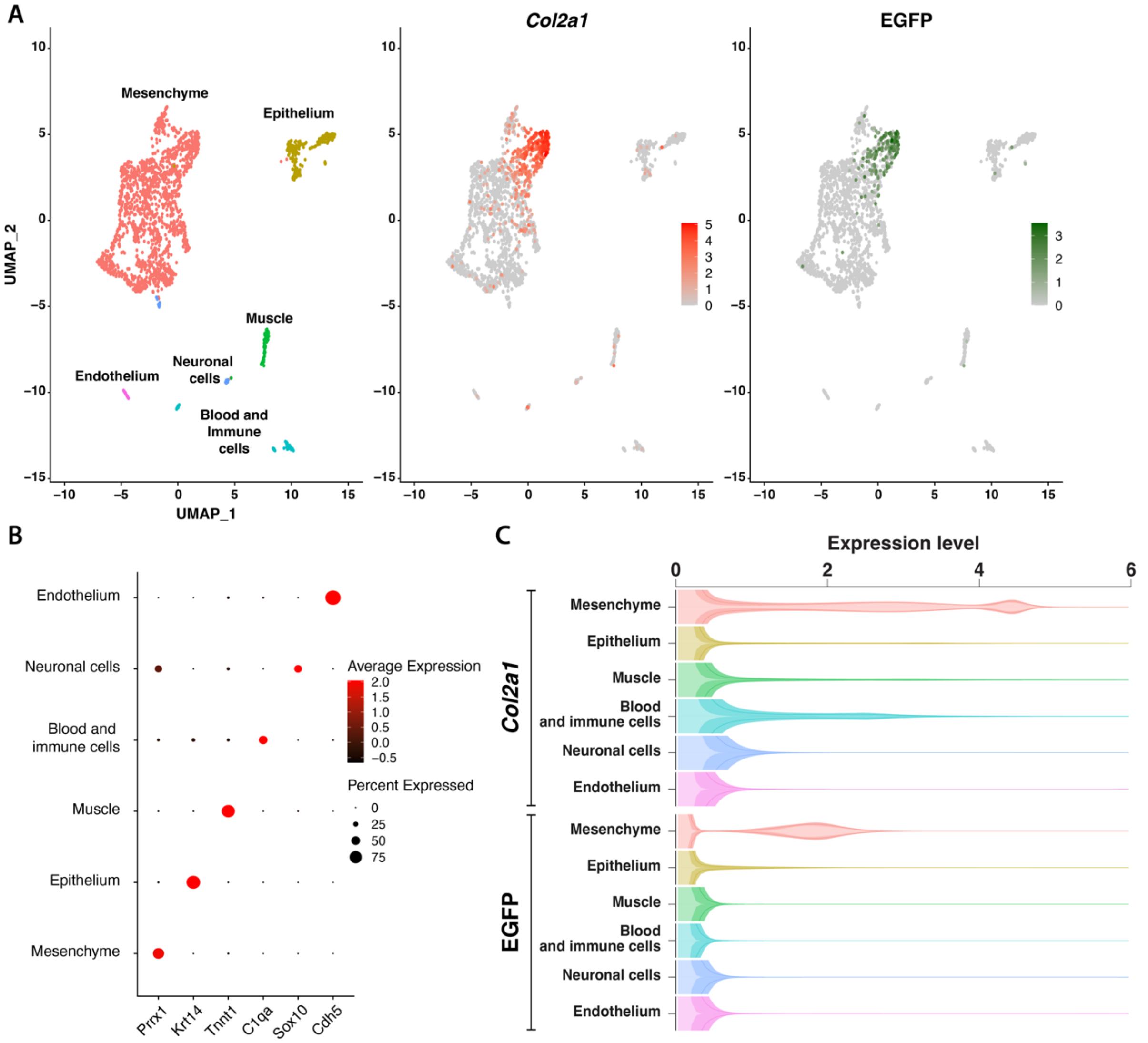


**Supplementary Figure 1:** Single-cell analysis of E14.5 *Col2a1*$^{EGFP/EGFP}$ limbs. **A.** Left panel: UMAP visualization of clustering of E14.5 limb cells. Middle: expression of *Col2a1* across the UMAP. Right: Expression of EGFP across the UMAP. **B.** Selected marker gene expression per cluster. **C.** Expression distribution of *Col2a1* and EGFP per limb cluster estimated by baredSC. Line shows mean and shaded area around the line indicates the 68% confidence interval. Density values, displayed on the y axis, were truncated at 0.3 and mirrored. Note the strong expression of both genes in the mesenchyme cluster.

### Supplementary Figure 2

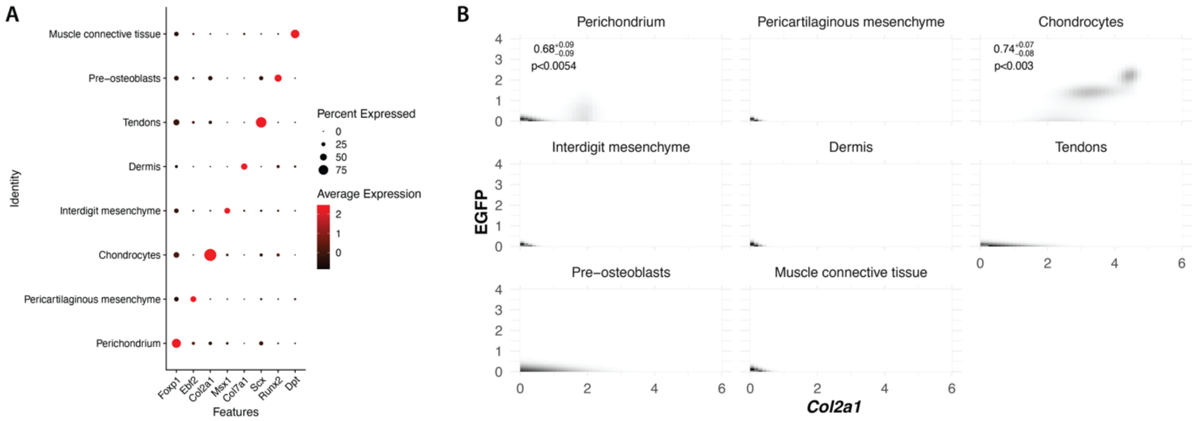


**Supplementary Figure 2: A.** Selected marker gene expression per mesenchymal cluster. **B.** Co-expression of *Col2a1* (x-axis) and EGFP (y-axis) in all limb mesenchymal clusters as determined by baredSC. For clusters with fraction of cells expressing *Col2a1* above 2, correlation coefficient is displayed with confidence interval and estimated p-value.

**Supplementary Figure 3**

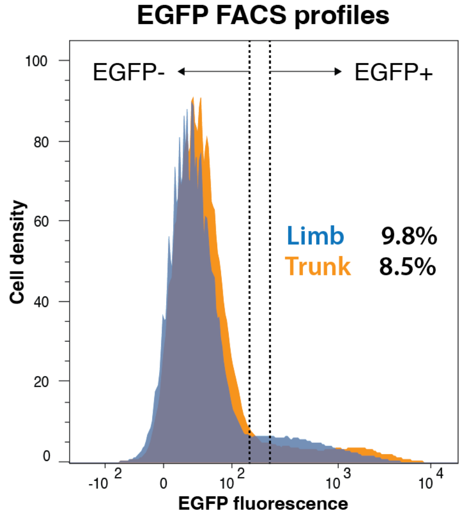


**Supplementary Figure 3:** FACS fluorescence profiles of E14.5 *Col2a1*$^{EGFP/EGFP}$ limbs and trunks.

### Supplementary Figure 4

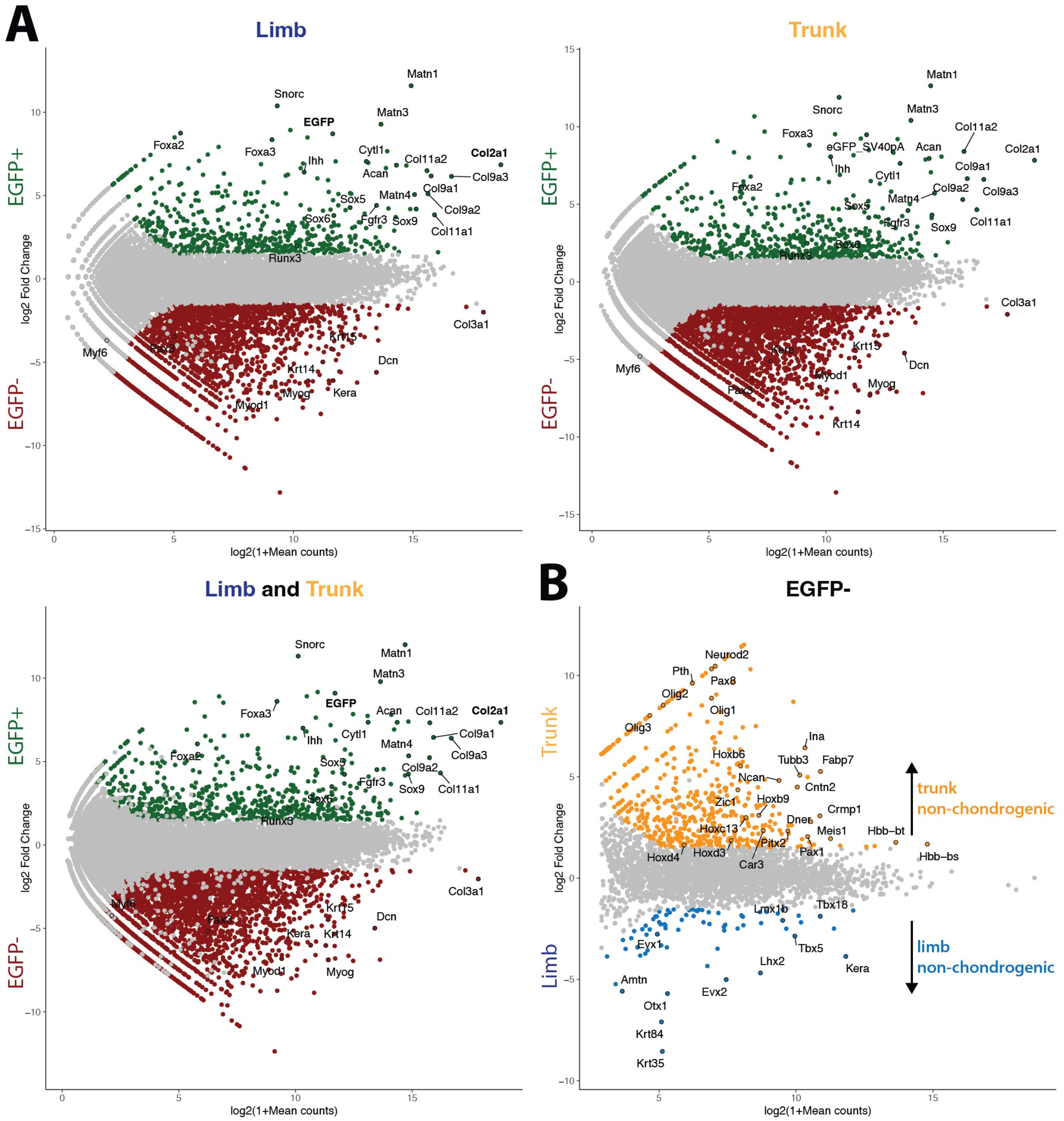


**Supplementary Figure 4: A.** Enriched genes in EGFP+ (green) and EGFP- (red) cell populations in limbs (top left), trunk (bottom left) and combined datasets (top right). **B.** Non-chondrogenic (EGFP-) marker genes with an expression preference in limb (blue) or trunk (orange).

**Supplementary Figure 5**

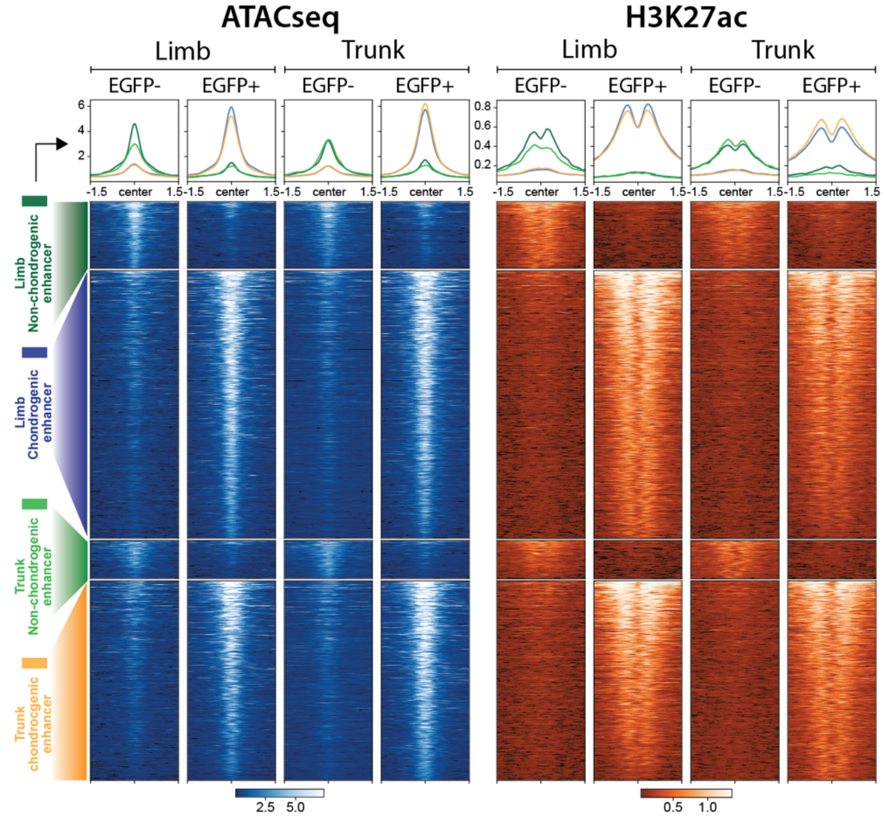


**Supplementary Figure 5:** Analyses of chondrogenic chromatin landscapes and enhancers. ATAC-seq and H3K27ac coverages over 3kb for limb and trunk chondrogenic and non-chondrogenic enhancers are centered at the corresponding merged ATAC-seq peaks located within a 75bp window. Enhancers might be present in multiple categories.

**Supplementary Figure 6**

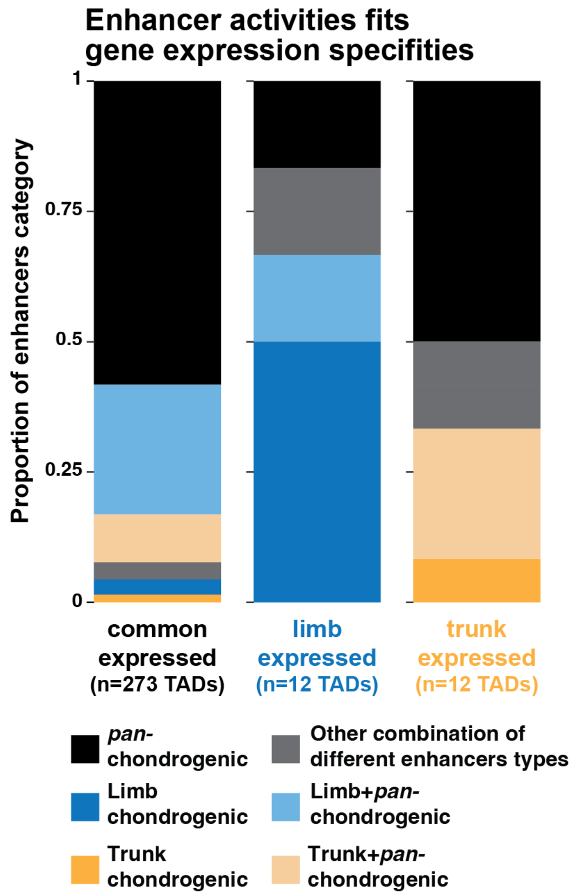


**Supplementary Figure 6:** Association between trunk- and limb- preference of chondrogenic protein coding genes and chondrogenic enhancers. Proportion of TADs containing the different enhancer types: only *pan*-chondrogenic enhancers (active in both limb and trunk chondrocytes), only limb enhancers (active preferentially in limb chondrocytes), only trunk enhancers (active preferentially in trunk chondrocytes) and combinations thereof split by TADs containing common-, limb-, or trunk-expressed genes. Note that the of proportion limb and trunk enhancers is increased in limb or trunk-associated TADs.

### Supplementary Figure 7

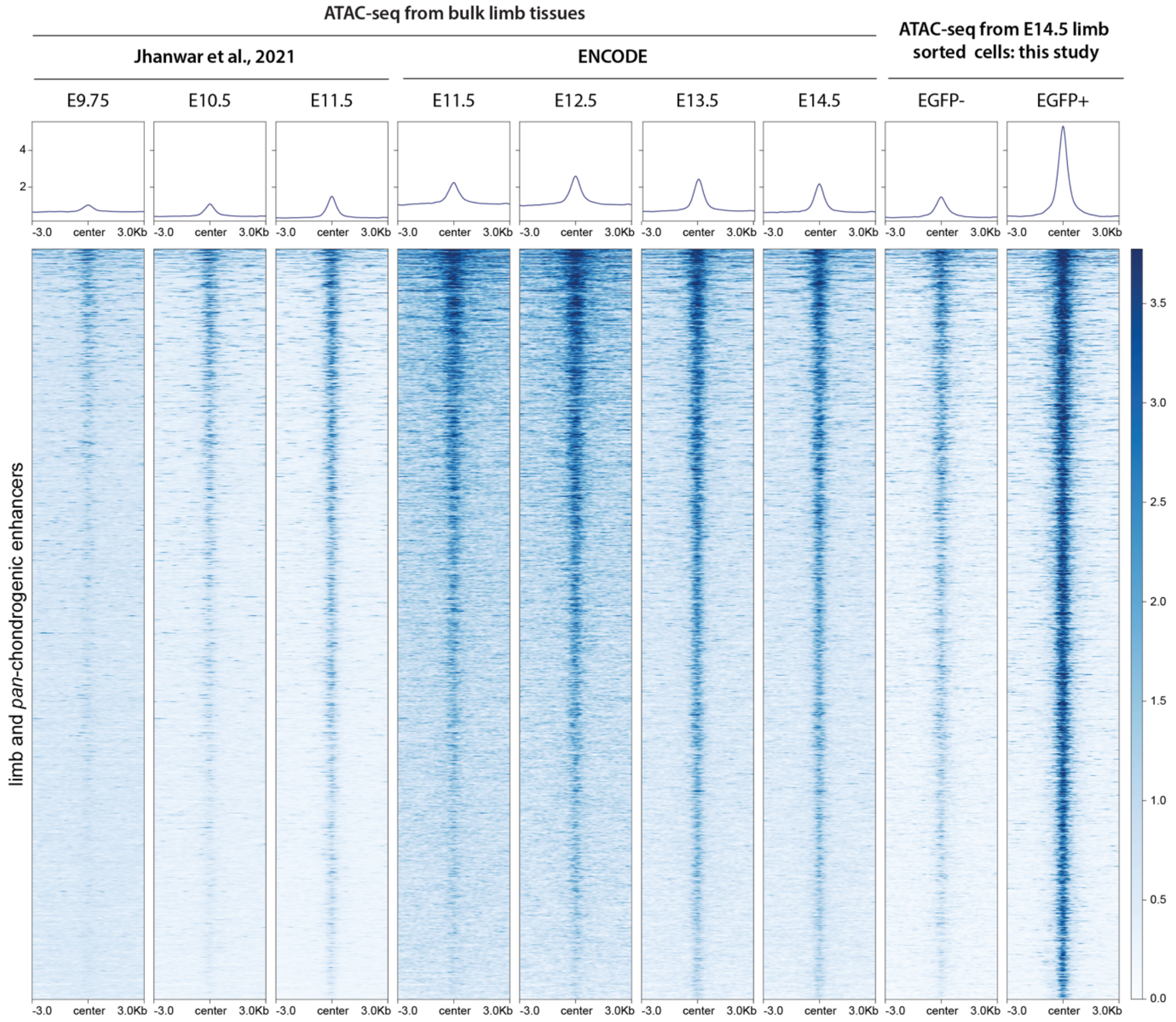


**Supplementary Figure 7:** ATAC-seq signal at chondrogenic enhancers over developmental time. At E9.75 only a few regions display signal while at E11.5 the maximum signal is obtained (to compare with E14.5 bulk). ATAC-seq coverages over 6kb are centered at the corresponding merged ATAC-seq peaks located within a 75bp window.

**Supplementary Figure 8**

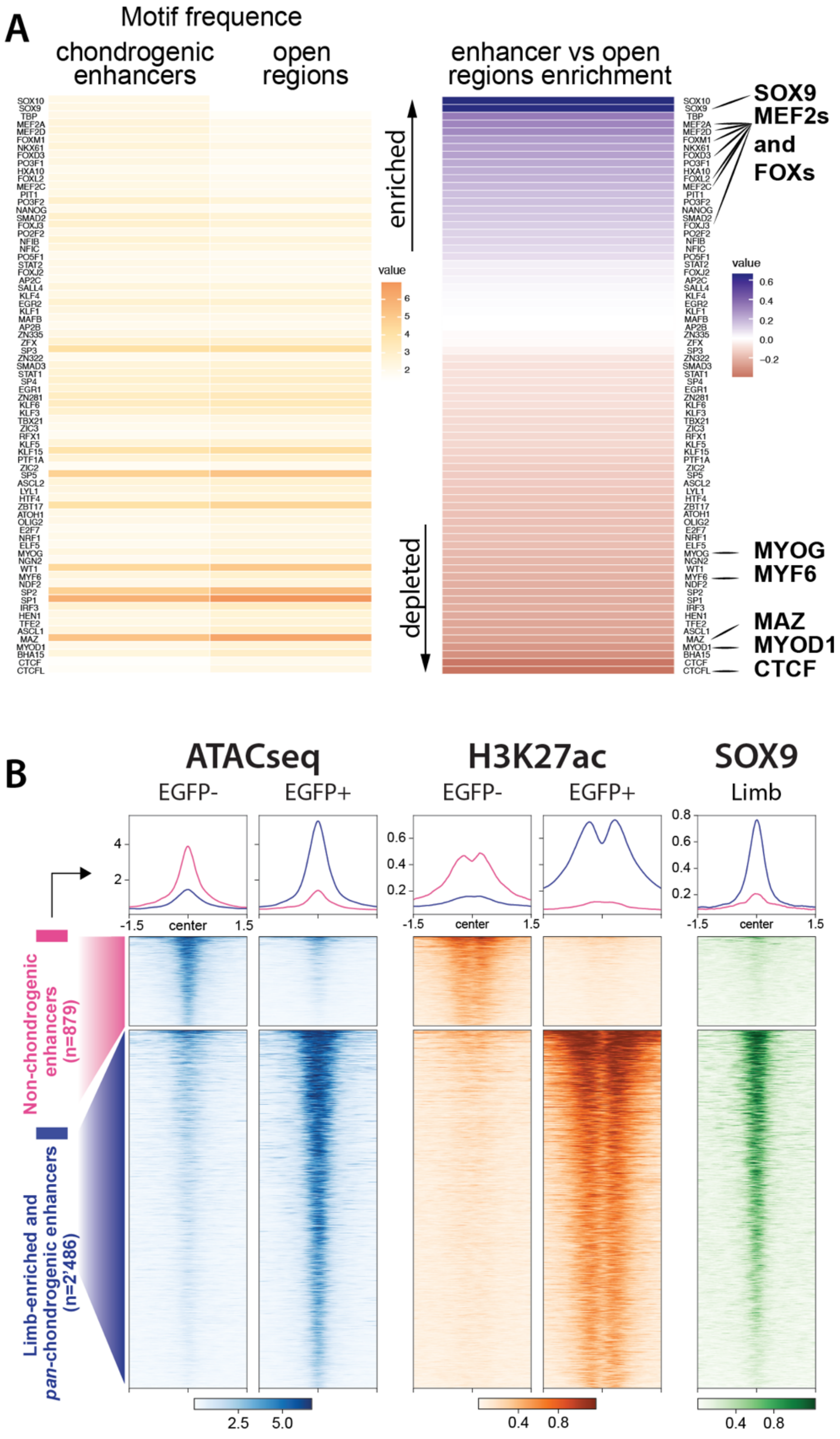


**Supplementary Figure 8:** Transcription factor binding at chondrogenic enhancers. **A.** Left: All enriched motifs between chondrogenic enhancers and "*inactive*" accessible regions. Right: ratio of motifs enrichment between chondrogenic enhancers and "*inactive*" accessible regions. **B.** Binding of *SOX9* at limb-enriched and *pan*-chondrogenic enhancers (n=2'486) compared to non-chondrogenic enhancers enrichment (n=879). Heatmaps are generated from the limb EGFP+ and EGFP- datasets. ATAC-seq, H3K27ac and SOX9 coverages over 3kb centered at the corresponding merged ATAC-seq peaks located within a 75bp window.

### Supplementary Figure 9

**A**

| Span_TAD_Castro_mm39 | size (bp) | chondrogenic genes | chondrogenic genes [n] | chondrogenic enhancers [n] |
|---|---|---|---|---|
| chr5:41207343-41967343 | 760000 | Nkx3-2 | 1 | 3 |
| chr10:77815834-86415864 | 8600030 | Aire, Chst11, Fstl3, Odf3l2, Slc41a2 | 5 | 18 |
| chr18:58093072-59213072 | 1120000 | | | 3 |
| chr13:47213476-49893476 | 2680000 | Barx1, Ecm2, Id4, Ninj1, Omd, Rnf144b | 6 | 13 |
| chr1:57999159-59119159 | 1120000 | | | |
| chr15:40993396-42793396 | 1800000 | | | |
| chr6:119426961-120466961 | 1040000 | B4galnt3, Ninj2 | 2 | 3 |
| chr11:111430826-113510826 | 2080000 | Sox9 | 1 | 33 |
| - | **19200030** | | **15** | **73** |

**B**

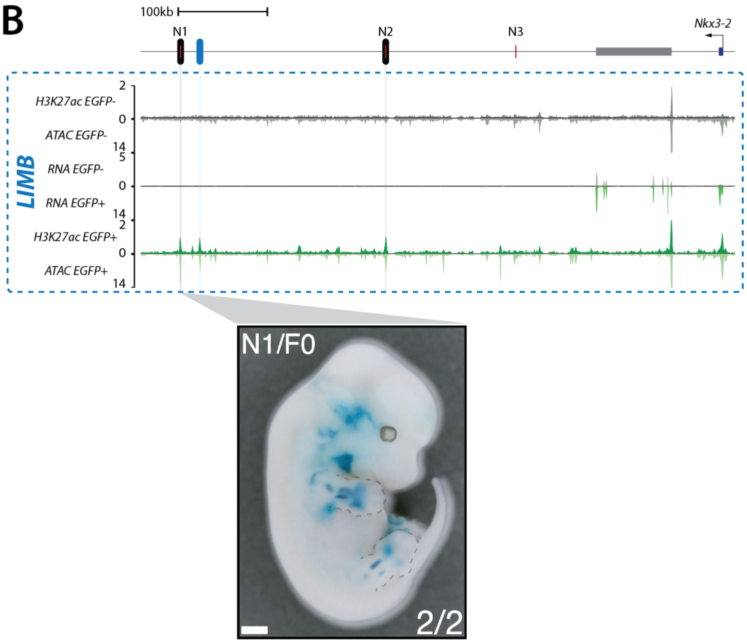


**Supplementary Figure 9: A.** Eight loci under selection were identified in Longshank mice, six of which contain numerous chondrogenic genes and enhancers (*40*). **B.** Distribution of RNA-seq, ATAC-seq and H3K27ac ChIP-seq normalized coverages in limb (blue box) and trunk (orange box) EGFP+ and EGFP- cells at the *Nkx3-2* loci. EGFP+ datasets are colored in green, EGFP- datasets in grey. RNA-seq and ATAC-seq coverages are an average of two replicates. Vertical bars highlight the genomic position of chondrogenic enhancers. At the *Nkx3-2* loci, we predicted 3 chondrogenic enhancers: 2 are categorized as *pan*-chondrogenic (black ovals) and 1 is limb-enriched (blue oval). 2 of the 3 chondrogenic enhancers have been identified as potentially affecting tibial length in longshank mice (N1 and N2). N3 was not predicted to be chondrogenic-specific. Photo was obtained from (*40*).

### Supplementary Figure 10

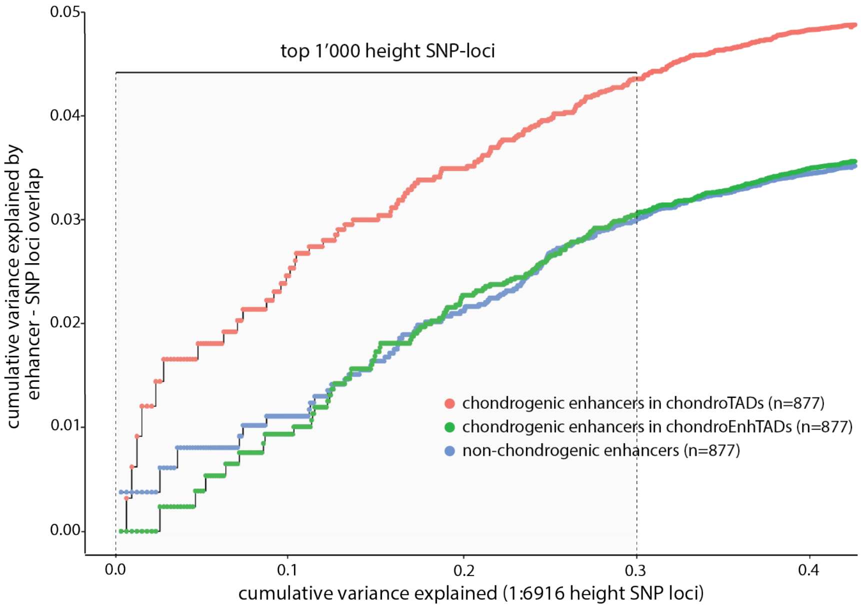


**Supplementary Figure 10:** Cumulative height variance explained by the 6'916 SNP-loci in mice (x-axis, regions sorted from highest to lowest variance) and the same variance explained by the enhancers overlapping the SNP-loci (y-axis). Enhancers on the X and Y chromosome were excluded from the analysis.

### Supplementary Figure 11

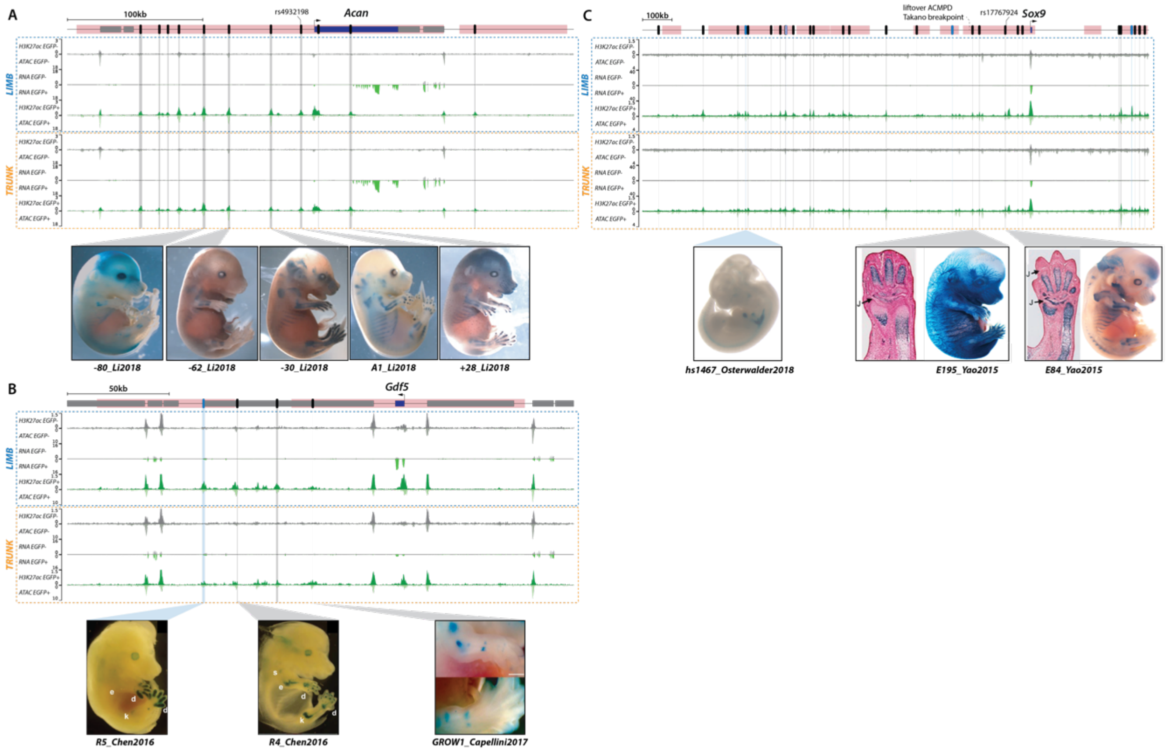


**Supplementary Figure 11: A-B-C** Distribution of RNA-seq, ATAC-seq and H3K27ac ChIP-seq normalized coverages in limb (blue box) and trunk (orange box) EGFP+ and EGFP- cells at the *Acan*, *Sox9* and *Gdf5* loci**.** Mouse homologous regions to Yengo height-associated GWS loci are highlighted in pink (*42*). EGFP+ datasets are colored in green, EGFP- datasets in grey. RNA-seq and ATAC-seq coverages are an average of two replicates. Vertical bars highlight the genomic position of chondrogenic enhancers. **A.** 12 chondrogenic enhancers are predicted at the *Acan* locus and categorized as *pan*-chondrogenic (black ovals). 6 out of the 12 enhancers have already been shown to drive chondrogenic expression of marker genes. Photos were obtained from (*43, 44*). 1 human SNP associated by Yengo et al. with height was identified in the enhancer A1 (*42*). **B.** 4 chondrogenic enhancers are predicted at the *Gdf5* locus: 3 are categorized as *pan*-chondrogenic (black ovals) and 1 is limb-enriched (blue oval). 3 out of the 4 enhancers have already been shown to drive chondrogenic expression of marker genes. Photos were obtained from (*45, 46*). Shoulder (s), elbow (e), knee (k) and digits (d) are identified on the R4 and R5 photos. **C.** 29 chondrogenic enhancers are predicted at the mouse *Sox9* locus, 25 are categorized as *pan*-chondrogenic (black ovals) and 4 are limb-enriched (blue ovals). 3 out of the 29 enhancers have already been shown to drive chondrogenic expression of marker genes. Photos of E195 and E84 were obtained from (*47*) and the VISTA Enhancer browser for hs1467 (*98*). Hs1467 is published by Osterwalder et al. (*6*). 1 human SNP associated by Yengo et al. with height was identified in the enhancer E84 (*42*). Carpal and phalangeal joints (J) are identified in the E84 and E195 sections. Below the human *SOX9* locus with lift over chondrogenic enhancers and the breakpoint involved in acampomelic CD (*60*).
